# Symmetry and similarity in eco-evo: advective environments, dispersal, and climate change

**DOI:** 10.1101/2025.08.26.672437

**Authors:** Ido Filin

## Abstract

I investigate effects of directional advective dispersal on eco-evolutionary invasion speeds, in either constant environmental gradients or under directional environmental change. I build upon a spatially-explicit phenotypic model that I have previously presented. The model incorporates both phenotypic sorting and gene swamping, and is used to study constant advective dispersal and trait-based habitat choice. Through analytical arguments on characteristic lengths and speeds, as well as by direct numerical solutions, I find that, in eco-evo scenarios, advective dispersal has two opposing effects on invasion fronts. In shallow environmental gradients, the demographic effect of simply adding to the steady-state invasion speed dominates. As gradients get steeper, asymmetric gene flow, due to advective dispersal, becomes increasingly stronger, pulling the invasion front in an opposite direction. Steep gradients cause front reversals, where species ranges, counter-intuitively, slide upstream. Symmetry between velocity of advective dispersal and velocity of climate change leads to adaptation lags, abundance profiles, and invasion speeds that depend only on the velocity difference. Through characteristic lengths and speeds, I also identify similarity between demographic and evolutionary averaging or tracking of environmental heterogeneity. I connect climate change research to demographic models of persistence in advective environments — the so-called drift paradox. I additionally derive dimensionless ratios that have potential in facilitating comparisons among different species, populations, traits and locales, and in guiding micro- and mesocosm experiments and space-for-time substitutions. As an aside, I demonstrate how numerical solution and simulation of complex eco-evo models can be accelerated with GPU programming.

## Introduction

There is an inherent symmetry between the velocity of climate change and the directional dispersal or flow that organisms experience in advective environments, such as rivers, propagule-carrying ocean currents and seed-dispersing winds. Though buried now in the older strata of an ever more recent literature, earlier theoretical work on population persistence under climate change (Potapov & Lewis 2004, Berestycki et al. 2009) originated in such persistence models for advective environments, namely, on the so-called drift paradox (Speirs & Gurney 2001, Pachepsky et al. 2005, Shanks & Eckert 2005, Lutscher et al. 2006, Nisbet et al. 2010, Vasilyeva & Lutscher 2012). In fact, when solving for conditions of persistence under climate change Potapov & Lewis (2004) applied the argument of “habitat motion as advection of biota” and, accordingly, transformed the initial climate-change formulation into a model of an advective environment.

There has also been much renewed interest in similar classic reaction-diffusion models of biological invasions (Skellam 1951, Shigesada & Kawasaki 1997, Murray 2002). Especially by recent studies that apply swarming microbes (Ben-Jacob et al. 1998) as microcosm models for eco-evolutionary (hereafter, *eco-evo*) dynamics of adaptation and range expansion (Gandhi et al. 2016, Deforet et al. 2017, Birzu et al. 2019, Erm & Phillips 2020).

Eighteen years ago, I presented (Filin 2007) a phenotypic reaction-transport (also reaction-diffusion, or reaction-advection-diffusion; Ishihara et al. 2020, Kumaran 2023) eco-evo model that combined gene swamping (Pease et al. 1989, Kirkpatrick & Barton 1997, Lenormand 2002), spatial sorting (Shine et al. 2011) and matching habitat choice (Edelaar et al. 2008), though the latter two terms were not yet invented. I then provided a systematic exploration of different forms of *advective dispersal* (also, directional, asymmetric, nonrandom, anisotropic, directed, biased, unidirectional, taxis, convection or drift; Gurney & Nisbet 1975, Nagylaki 1978, van den Bosch et al. 1990, Farnsworth & Beecham 1999, Levine 2003, Hare et al. 2005, Filin 2007, Nisbet et al. 2010, Sorte 2013, Potts & Schlägel 2020, Hancock et al. 2024), and how they affect eco-evo dynamics and range expansions. Here, I recap and elaborate on my approach and some of my results, present them within current eco-evo context, and provide new analyses, to demonstrate how the study of advective dispersal highlights the nuances of eco-evo dynamics.

Two classic results of spatially-explicit reaction-diffusion population models in ecology are critical domain (or patch) size and constant steady-state invasion speed (Skellam 1951, Kierstead & Slobodkin 1953, Okubo 1984, Shigesada & Kawasaki 1997, Potapov & Lewis 2004, Nisbet et al. 2010). Critical domain size defines a demographic characteristic length, over which spatial variation in population (per-capita) growth (e.g., spatial variation in recruitment; Nisbet et al. 2010) is averaged. A population may not be able to persist in an otherwise favorable source patch, if surrounding sinks dominate in this spatial averaging. In other words, critical domain size defines the finest spatial resolution of habitat heterogeneity, to which populations can respond, switching from averaging spatial heterogeneity to tracking it (Roughgarden 1974, Anderson et al. 2005, Nisbet et al. 2010). When random dispersal is denoted by a diffusion coefficient, *D*, and per-capita growth rate by *r*, the critical patch size is proportional to 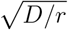 (Kierstead & Slobodkin 1953, Nisbet et al. 2010).

In invasion scenarios, the quantity 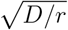 also defines the shape of the advancing front, i.e., the spillover or decay of population density with distance (Gandhi et al. 2016). The speed at which this invasion front propagates, through population growth and random dispersal, is 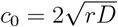 (Skellam 1951, Shigesada & Kawasaki 1997). This is the steady-state invasion speed of pulled waves (Birzu et al. 2019). I will refer to *c*_0_ as the *Fisher velocity* (also, Fisher wave speed; Fisher 1937, Nisbet et al. 2010, Gandhi et al. 2016, Birzu et al. 2019). It is a baseline invasion wave speed for homogeneous environments, negatively density-dependent growth (i.e., no Allee effect), and only diffusive random dispersal.

In eco-evo models, however, evolutionary dynamics, such as of the mean phenotype or trait distribution across space, also introduces characteristic dimensions of its own. Specifically, for mean trait, the evolutionary characteristic length, hereafter evolutionary *response length* (borrowing from the purely demographic model of Anderson et al. 2005), is given by 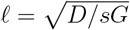 (May et al. 1975, Endler 1977, Slatkin 1978, Lande 1982b, 1991, Barton 1999, Kruuk et al. 1999, Day 2000, Filin 2007), where *s* is a measure of the strength of stabilizing selection towards a spatially varying local optimum, and *G* is the trait’s additive genetic variance within local populations. Like the ecological critical domain size, the evolutionary response length defines averaging by random dispersal of spatially varying selection (Lande 1982b). Consequently, local mean phenotype is a weighted average of spatially varying optima, determined by an averaging kernel that decays with distance according to *ℓ* (Slatkin 1978, Lande 1982b, Filin 2007).

There is also an evolutionary characteristic speed, analogous to *c*_0_ of demographic invasion dynamics, 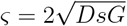 (clearly related to Fisher‘s 1937 original formulation for spread of advantageous mutations; compare also with the critical speed of Kirkpatrick & Peischl 2013, for evolutionary rescue in single-locus models). I will refer to it here as the *characteristic speed*. In the context of this work, this measure does not describe spatial spread *per se*. But, rather, it defines a velocity scale, by which other velocities are weighed to influence mean trait values.

Of course, the heart of eco-evo is the coupling of the two types of dynamics — demographic and evolutionary. Evolution affects demography via local maladaptation, deviation of mean trait from local optimum, which depresses mean fitness and per-capita growth (Kirkpatrick & Barton 1997, García-Ramos & Rodriguez 2002, Barton 2024). In the opposite direction, the effect of demography on evolution is, traditionally, through mixing by random dispersal (or migration), and through gene swamping — the tendency of the mean trait to deviate from the local optimum towards optima of more populous localities, due to the relatively stronger incoming (asymmetric) gene flow from those localities (Pease et al. 1989, Kirkpatrick & Barton 1997, Lenormand 2002). Gene swamping, therefore, distorts the evolutionary averaging kernel, extending it upstream, while contracting downstream (Slatkin 1978, Filin 2007).

From a modeling perspective, gene swamping introduces a directional *advection* term to evolutionary dynamics (asymmetric gene flow), while no advection is present in demographic dynamics, if dispersal is random and symmetric. Advection as asymmetric gene flow has been previously explored in models of allele-frequency clines (May et al. 1975, Nagylaki 1978) and spatially-varying selection-migration models of quantitative traits (Slatkin 1978, Hare et al. 2005). Those studies, however, do not include explicit demographic dynamics, assuming a perscribed spatial profile of population density (usually uniform; but see Hare et al. 2005). In purely demographic models, directional advective dispersal, denoted by advection velocity *u*, adds to the rate of spatial spread, resulting in a combined invasion speed, *c* = *c*_0_ + *u* (van den Bosch et al. 1990, Shigesada & Kawasaki 1997). Advection velocity also affects conditions for persistence of populations in such advective environments as rivers and streams, wind- and waterborne seed dispersal, or larval dispersal of marine invertebrates — the so-called drift paradox (Speirs & Gurney 2001; drift in the sense of advective flow). Conditions for persistence depend on velocity of advective dispersal, in conjunction with the critical domain size, 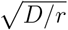, and the Fisher velocity, *c*_0_ (Speirs & Gurney 2001, Nisbet et al. 2010). The motivation of my investigation is to explore how such advective dispersal also modulates the full two-way feedback between evolution and demography in eco-evo scenarios, to affect response lengths, invasion speeds and range pinning, in either constant environmental gradients or under directional environmental change.

## Methods

The notation I adopt here is common in physics and theoretical ecology (Gurney & Nisbet 1975, Shigesada & Kawasaki 1997, Okubo & Levin 2001, Ishihara et al. 2020). Somewhat different notation is used in population genetic models (Pease et al. 1989, Kirkpatrick & Barton 1997, García-Ramos & Rodriguez 2002, Polechová et al. 2009). In the SI, I provide a brief guide for the perplexed on notation.

In the Appendix, I derive (following Filin 2007) a spatially-explicit phenotypic reaction-transport model for population density, *n*(***x***, *t*), and frequency distribution of trait values within local populations, *f* (*z*, ***x***, *t*). Random transport is denoted by a diffusion coefficient, *D*, and a directional component of dispersal (advective, or convective, transport) is denoted by an advection velocity vector ***u***. In general, both diffusive and advective transport may depend on space, ***x***, time, *t*, trait or phenotypic value, *z*, and population density, *n*. This leads to relative-fitness-like terms that depend on relative random dispersal, 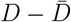, and relative advective dispersal, 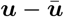, demonstrating the equivalence of selection and spatial sorting (Shine et al. 2011, Phillips & Perkins 2019).

In addition to sorting, the general model also includes the effects of gene flow asymmetry, or gene swamping (Pease et al. 1989, Kirkpatrick & Barton 1997, Lenormand 2002), summarized through the vector

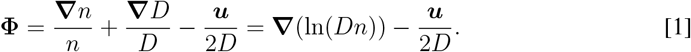

The operator **∇** (gradient operator) represents spatial derivatives, i.e., with respect to spatial position, ***x***. In this context, it describes spatial variation in local population density and in rates of diffusive transport (spatial variation in random dispersal). The gene flow asymmetry vector, **Φ**, captures the equivalence of spatial gradients in relative population density, spatial variation in random dispersal rates, and directional advective transport. This equivalence has been partly shown by Fife (1979) and Nagylaki (1989) in allele-frequency cline models, and by Gurney & Nisbet (1975) in a purely demographic model.

The simplest form of advective dispersal is a fixed velocity, ***u*** = *const*. It can be interpreted as the effect of a prevailing wind direction, flow in a river, or as asymmetry in the spatial distribution of dispersers or propagules (asymmetric dispersal kernels; Slatkin 1978, Okubo & Levin 1989, Levine 2003). Similarly, directional temporal environmental change, hereafter *climate change*, can be described by a linearly time-varying environmental gradient, given by *θ*(***x***, *t*) = ***b* ·** (***x*** − ***u***_*θ*_*t*), where *θ* is the spatially and temporally varying optimal trait value, ***u***_*θ*_ is the velocity of climate change (Pease et al. 1989, Loarie et al. 2009), and ***b*** represents the slope of the environmental gradient, measuring how steeply selective forces vary across space (Kirkpatrick & Barton 1997, García-Ramos & Rodriguez 2002). (This notation is equivalent to *θ* = *bx* − *kt* of Bürger & Lynch 1995 and Lande & Shannon 1996, in one-dimensional space, where *k* denotes temporal rate of change in local conditions; *k* = *bu*_*θ*_.)

I study the eco-evolutionary effects of advective transport through the dynamics of spatiotemporally varying population density, *n*(***x***, *t*), and mean trait, 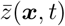. This is achieved by substituting particular forms of ***u***(***x***, *z*) (i.e., possibly trait-dependent, as in the next paragraph) in the general model for population density and trait distribution (Eqs.[12] and [13] in Appendix), and deriving expressions for *∂n*(***x***, *t*)/*∂t* and for dynamics of local (mal)adaptation, *∂a*(***x***, *t*)/*∂t* (Eqs.[14]–[21] in Appendix), where 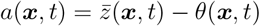 (also called adaptation lag or biotic lag; Lande & Shannon 1996, Lovell et al. 2023).

Beyond a fixed velocity, there are many other forms of advective dispersal that can be explored. Here, I only present phenotype-dependent (matching) habitat choice (Filin 2007, Edelaar et al. 2008). (Additional forms were explored in Filin 2007; see also Hare et al. 2005.) In an analogous manner to evolving populations climbing up fitness gradients in (multivariate) trait space (Lande 1982a), matching habitat choice can be viewed as advective dispersal up spatial fitness gradients, *u*(*x, z*) = *χ***∇***w*(*x, z*), where *w*, here, denotes fitness (Filin 2007; Appendix). The proportionality coefficient, *χ*, was adopted from models for chemotaxis (Keller & Segel 1971), where it is called chemotactic mobility and measures the sensitivity of movements of cells to concentration gradients of nutrients. In the context of habitat choice, this parameter translates fitness differences among neighboring environments to rates of advective dispersal. It has dimensions of length squared. Therefore, given an environmental gradient of steepness ***b***, the expression 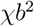 has dimensions of trait variance.

Environmental gradients, where optimum trait value varies across space, are commonly associated with quadratic (or, in discrete time, Gaussian) fitness functions, representing stabilizing selection towards the spatiotemporally varying optimum, with selection intensity measured by parameter *s* (García-Ramos & Rodriguez 2002). Given a linear environmental gradient, *θ*(***x***, *t*) = ***b* ·** (***x*** − ***u***_*θ*_*t*), matching habitat choice translates, in this case, to mean advective dispersal, 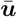, and a sorting term, 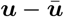, given by

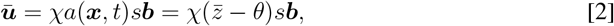

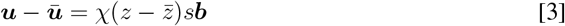

(Filin 2007; Appendix). These expressions can, again, be substituted in the general dynamics equations for population density and trait distribution, to produce the required expressions for *∂n*(***x***, *t*)/*∂t* and *∂a*(***x***, *t*)/*∂t* (see Appendix).

Effects of diffusive and advective transport on dynamics of *n* and *a* (or 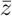) were derived under fairly general conditions, without assuming particular forms of population growth, trait distribution and selection (except for quadratic fitness function in the case of habitat choice). For the purpose of getting closed-form analytical expressions and for the numerical examples (Fig. 1), trait variance will be taken as a fixed parameter, and stabilizing selection in the dynamics of mean trait, or maladaptation, will follow the typical formulation of 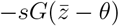, or −*sGa* where *G* is the additive genetic variance (Slatkin 1978, Bürger & Lynch 1995, García-Ramos & Rodriguez 2002). In numerical solutions, I will additionally make the simplifying assumptions of one-dimensional space, ***x*** = *x* (dropping the vector notation), and logistic population growth (Eqs.[18]–[21] in Appendix).

**Figure 1:**
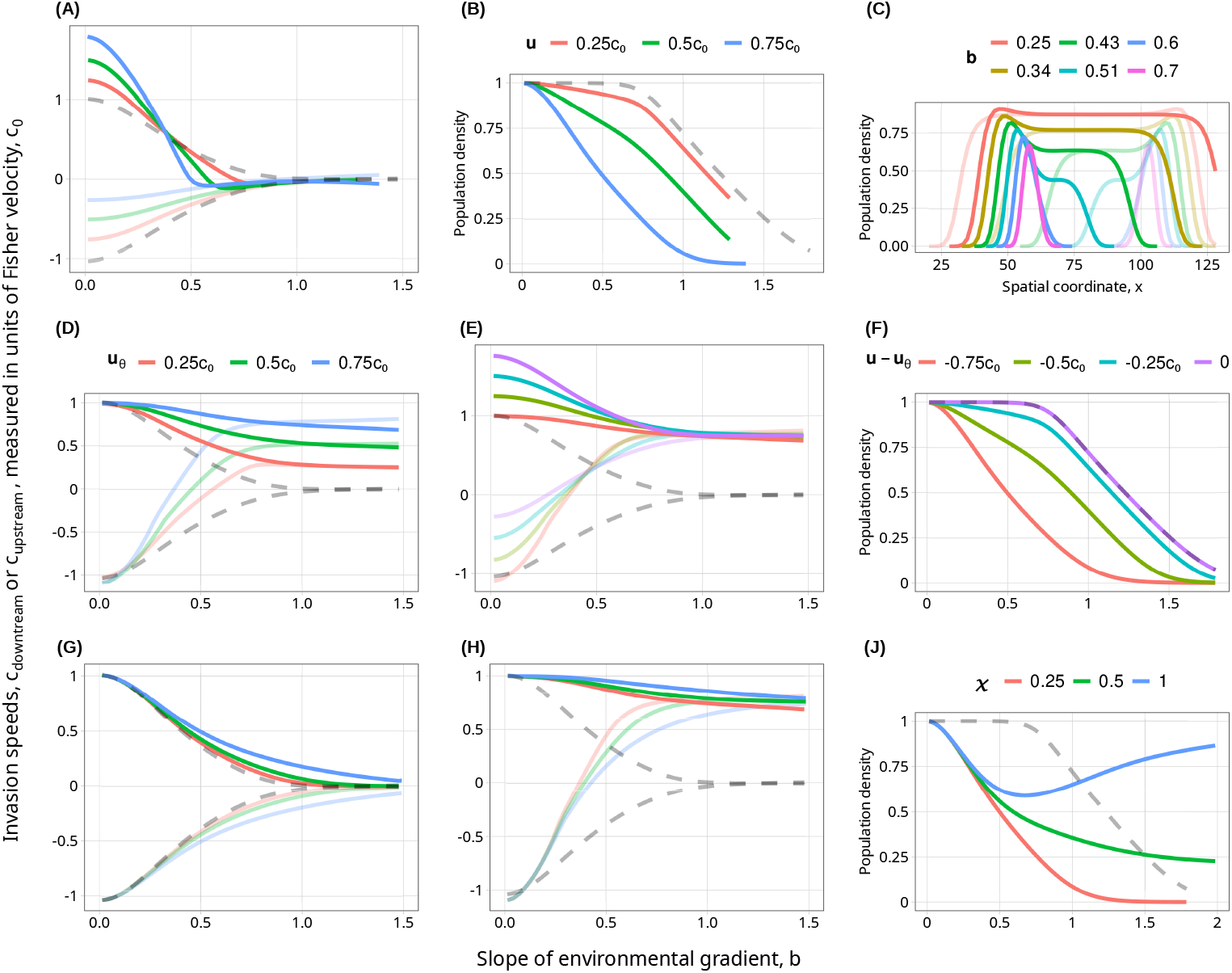
Invasion speeds of both downstream (solid lines) and upstream (transparent lines) wave fronts and population densities, as functions of slope of environmental gradient, *b* (except C). As the velocity of advective dispersal, *u*, increases (A; legend in B), invasion speeds shift downstream. Dashed line represents the case of no advective dispersal and no climate change (*u* = *u*_*θ*_ = 0), i.e., purely symmetric random dispersal in a constant environmental gradient (invasion speeds in this case are in agreement with Figure 6 of Filin et al. 2008 for genetic variance parameter *A* = 0.5 and logistic density-dependence). Clearly, for the purely demographic case (*b* = 0), *c* = *c*_0_ + *u*, as expected (Shigesada & Kawasaki 1997). For steeper gradients, invasion speeds decline in both downstream and upstream fronts (as expected for evolutionary speeds of invasion; García-Ramos & Rodriguez 2002). Range pinning, where curves of downstream and upstream velocities converge, occurs at increasingly shallower gradients (lower values of *b*), as advective dispersal, *u*, increases. Final maximum population density (B) also decreases with both steepness of environmental gradient, *b* and velocity of advective dispersal, *u*. Filin (2007), in fact, derived a condition for persistence in the environmental gradient 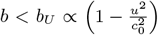 (cf. Speirs & Gurney 2001, Potapov & Lewis 2004, Nisbet et al. 2010, in purely demographic scenarios of advective flow). For velocity of climate change, *u*_*θ*_, effects are overall similar (D), though in this case, invasion speeds decline towards *u*_*θ*_, not 0 (compare to A). The symmetry is also evident when comparing spatial abundance profiles (C). Profiles for *u* = 0.5*c*_0_ (solid lines) and profiles for *u*_*θ*_ = 0.5*c*_0_ (transparent lines) are mirror images of each other, around the midpoint of initial introduction. Thus, with advective dispersal, densities shift upstream, due to pulling by gene swamping. With velocity of climate change they are higher downstream (cf. Possingham & Roughgarden 1990, Gaylord & Gaines 2000, Berestycki et al. 2009), towards shifting environments priorly adapted to. Such non-abundant-center density profiles suggest that, in eco-evo scenarios, even without stochasticity, nonequilibrium processes, in this case, ongoing spatial spread and (mal)adaptation, produce maximal densities closer to range margins (cf. Gaylord & Gaines 2000, Sagarin & Gaines 2002, Faurby & Araújo 2018, Dallas et al. 2020). With both advective dispersal and climate change (E; legend in F), dynamics depends on the velocity difference *u* − *u*_*θ*_. When *u* matches *u*_*θ*_, one clearly regains the symmetric random dispersal case (dashed line), only shifted downstream by *u*_*θ*_ (here *u*_*θ*_ = 0.75*c*_0_). Symmetry is also evident in final maximum densities (F; compare with B). Finally, G, H, J (common legend in J) demonstrate the added effect of matching habitat choice, through the parameter *χ*. Clearly, population density (persistence) is enhanced, invasion speeds are higher (Filin 2007), and range pinning requires steeper environmental gradients, as habitat choice gets stronger (cf. Pellerin et al. 2019). *u* = *u*_*θ*_ = 0 in G. *u*_*θ*_ = 0.75*c*_0_ in H, J (compare with D).

Finally, numerical solution of the eco-evolutionary dynamics equations was implemented with GPGPU — general purpose computing on graphics processing units — and run on a personal laptop computer. Programming was done in JavaScript, WebGL and GLSL (OpenGL shader language). This allowed me to write, compile and run programs on the vastly parallel GPU, and to utilize and access GPU memory (video memory) directly. Such hardware acceleration of numerical procedures allowed for quick software development iterations and exploration of a wide chunk of parameter space. I used the classic explicit finite-difference method for parabolic equations (Smith 1985), which, although lacking in sophistication, easily lends itself to parallelization. As in Filin (2007), I numerically solved for log-transformed population densities, ln(*n*), rather than *n* directly. More details on numerical procedures in SI.

## Results

With advective dispersal at velocity ***u***, an equilibrium solution of uniform population density across the entire environmental gradient is characterized by a constant level of local maladaptation, *a* = −***u* · *b****/sG*. This expression is the mirror image of the lag in adaptation due to climate change at velocity ***u***_*θ*_ (Lande & Shannon 1996, Polechová et al. 2009), resulting in the combined expression

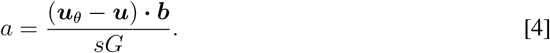

This symmetry between velocity of climate change, ***u***_*θ*_ and velocity of advective dispersal, ***u***, suggests a possible interesting direction for space-for-time substitutions (Lovell et al. 2023) in climate change research. Namely, studying populations that have historically adapted under conditions of advective dispersal (Sorte 2013), as an analogy for future adaptation (or failure to adapt) under climate change. It also points at advective dispersal as a possible adaptive response to negative effects of directional environmental change, as demonstrated in Fig. 1E,F. That can be grasped intuitively because, as optimal environments shift downstream (e.g., poleward), advective dispersal helps to restore a match between incoming dispersers and local environment.

Symmetry is also evident when exploring so-called evolutionary speeds of species invasions (García-Ramos & Rodriguez 2002, Filin et al. 2008). For shallow environmental gradients (small *b*), an approximate expression for invasion speed, *c*, relative to velocity of climate change, *u*_*θ*_, is

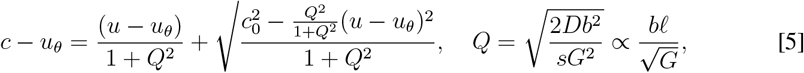

combining random dispersal (*D*), advective dispersal (*u*), velocity of climate change (*u*_*θ*_) and effects of environmental heterogeneity (*b*) (see Appendix for derivation). This relative invasion speed clearly depends, as for adaptation lag (Eq.[4]), only on the velocity difference *u* − *u*_*θ*_.

The dimensionless quantity *Q* is the change in environmental conditions (optimum trait value) over a response length, measured in additive genetic standard deviations. In homogeneous environments (*b* = *Q* = 0), Eq.[5] becomes the purely demographic invasion speed *c*_0_ + *u*. For no climate change and no advective dispersal (*u* = *u*_*θ*_ = 0), Eq.[5] reduces to a previously presented expression for “critical speed” (Miller 2019).

I will refer from now on to the direction opposite to *u* as *upstream*, and the direction of *u* as *downstream* (Levine 2003, Sorte 2013). A systematic computational exploration, from shallow to steep environmental gradients, reveals that, as gradients get steeper, there can be reversal of range expansion, causing slowly expanding and pinned ranges (Kirkpatrick & Barton 1997) to, counter-intuitively, slide upstream. This can be understood by considering the balance of forces operating on range expansion in eco-evo scenarios. Demographic invasion models show that invasion wave fronts are pulled forward (Shigesada & Kawasaki 1997, Gandhi et al. 2016) by population growth and dispersal at the advancing front. However, in eco-evo models asymmetric gene flow, in this case

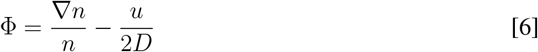

(Eq.[1] with *D* = *const* and one-dimensional space, ∇ = *∂*/*∂x*), causes the evolutionary averaging kernel to become asymmetric and biased upstream. This induces maladaptation at the advancing front, depressing growth rates and slowing invasions. Directional advective dispersal, *u >* 0, adds to this upstream bias, on top of gene swamping by density gradients, as Eq.[6] demonstrates.

The effect of advective dispersal on invasion speed is, therefore, on the one hand, to add to the downstream invasion speed — the purely demographic effect, *c* = *c*_0_ + *u* (Shigesada & Kawasaki 1997). On the other hand, advective dispersal increases the upstream “pull” of gene swamping, dragging the species range in an opposite direction. The balance of these opposing forces is dependent on the steepness of the environmental gradient, *b* (Fig. 1), and on the magnitude of upstream pulling by gene swamping. This upstream pulling is measured by a dimensionless quantity of gene flow asymmetry,

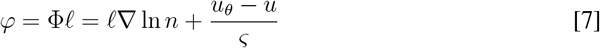

(Appendix). (For uniform population densities, ∇ ln *n* = 0, and no climate change, *u*_*θ*_ = 0, this expression is equivalent to the asymmetry parameter of May et al. 1975, and to 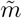 of Nagylaki 1978, for single-locus allele-frequency clines.)

For uniform population density this expression is simply (*u*_*θ*_ − *u*)*/ς*. For population densities that decrease exponentially with distance, *n* ∝ *e*^−*x/λ*^ (as in demographic invasion fronts; Murray 2002; *λ* here represents decay length; Gandhi et al. 2016, Deforet et al. 2017), gene flow asymmetry is given by *φ* = −*ℓ/λ* − (*u* − *u*_*θ*_)*/ς*, namely, the relative drop in population density over a response length minus the advective dispersal velocity, relative to velocity of climate change, scaled by the characteristic speed. This dimensionless measure of gene flow asymmetry can be either positive or negative, determining the direction and magnitude of upstream distortion in the evolutionary averaging kernel. The averaging distance expands upstream to 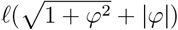, and contracts downstream to 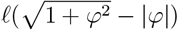 (Slatkin 1978, Filin 2007; Appendix). The total geographic extent of evolutionary averaging also widens, from 2*ℓ* to 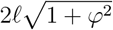, further precluding local adaptation and geographic differentiation.

A third point of symmetry between velocity of environmental change, *u*_*θ*_, and advective dispersal, *u*, is in spatial population density profiles during invasions. As Fig. 1C shows, climate change induces high population density close to the downstream edge of the expanding range, while advective dispersal is associated with highest densities near the upstream edge. In either case, maximum population density is closer to the range’s edges than to its center. However, when advective dispersal matches the velocity of climate change, *u* = *u*_*θ*_, symmetrically centered density profiles are regained (not shown).

Finally, trait-dependent habitat choice is simultaneously a form of advective dispersal (Eqs.[2] and [3]) and a form of spatial sorting (Shine et al. 2011, Phillips & Perkins 2019). Individuals within each local population disperse at varying rates and in different directions towards environments that better match their trait value. Intuitively, therefore, one expects such matching habitat choice (Edelaar et al. 2008) to reduce local maladaptation, matching mean trait values of populations to local optima and enhancing phenotypic differentiation across space. The response length accordingly shrinks to 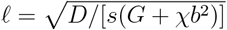, and the characteristic speed accelerates to 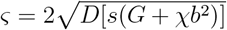 (Filin 2007; Appendix). Shorter response lengths translate to better tracking of spatially varying selection by local populations.

A shorter response length also translates to faster invasions (*Q* is decreased in Eq.[5]), as supported by numerical solutions (Fig. 1G,H). Higher characteristic speeds, similarly, decrease the upstream pull of gene swamping (*φ* is smaller; Eq.[7]). The averaging kernel is more symmetric and equilibrium levels of maladaption are lower (given now by *a* = (*u*_*θ*_ − *u*)*b/*(*sG* + *sχb*^2^); compare with Eq.[4]), enhancing overall persistence of the species in the environmental gradient (Fig. 1J). These expressions for characteristic length and speed, and adaptation lag, under trait-dependent habitat choice, clearly demonstrate that, in an eco-evo context, the role of such matching habitat choice, or spatial sorting, is equivalent to that of additive genetic variance, in promoting local adaptation. The effect manifests itself, though, only in moderate to steep environmental gradients (Fig. 1G,H,J), as it is proportional to *b*^2^.

## Discussion

A decade ago, Sorte (2013) discussed unique challenges of climate change to advective (flow) environments and to the species that depend on them, such as marine fishes and invertebrates, wind-dispersed plants, and lotic systems (Müller 1982, Nathan et al. 2011, Bertola et al. 2020). Sorte identified the direction of flow, relative to that of of climate change, as a key parameter in the future persistence of such populations. In this study, I have quantified this argument, through dependence of maladaptation (or biotic lag; Lovell et al. 2023), invasion speeds and spatial abundance profiles on the velocity difference, ***u*** − ***u***_*θ*_.

Population genetics often deals with questions of adaptation and persistence in terms of loads (Barton 2024), i.e., negative effects on fitness — genetic load (Crow & Kimura 1963), lag load (Lande & Shannon 1996, Polechová et al. 2009), expansion load (Peischl et al. 2013), and so on. However, in a spatial context, where dispersal influences a multitude of coupled demographic and evolutionary processes (mixing, averaging, swamping, sorting, etc.), there is value in looking at characteristic lengths and velocities as well. This approach has been productive in demographic models of spatial spread, of averaging and tracking of environmental heterogeneity, and of population persistence (Roughgarden 1974, Shigesada et al. 1986, Shigesada & Kawasaki 1997, Speirs & Gurney 2001, Potapov & Lewis 2004, Anderson et al. 2005, Berestycki et al. 2009, Nisbet et al. 2010), and more recently, in experimental microbial models for range expansion and invasion dynamics (Gandhi et al. 2016, Deforet et al. 2017, Birzu et al. 2019). The characteristic (response) length has also featured prominently in models of sexual selection and geographic variation in female mating preference (Lande 1982b, Day 2000), and in studies of allele-frequency clines (Slatkin 1973, May et al. 1975, Slatkin 1978, Nagylaki 1978, Barton 1999). Wave speeds have, similarly, been studied for selective sweeps (Fisher 1937, Barton et al. 2013) and evolutionary rescue (Kirkpatrick & Peischl 2013).

That is the second symmetry, more precisely similarity, that I have identified in this study. Namely, the dynamics of tracking and averaging (Roughgarden 1974, Anderson et al. 2005, Diehl et al. 2008) of environmental heterogeneity in both demographic and evolutionary dynamics, albeit with different characteristic length and speed scales. A more solid appreciation of this similarity in coupled eco-evo tracking and averaging, of transitions between the two modes, and of their modulation by gene swamping, and other eco-evo processes, should be sought.

Moreover, by identifying characteristic times, lengths and velocities, it is possible to make educated guesses about how additional processes and mechanisms may influence eco-evo dynamics, without initially having to solve complex models. Here, I demonstrated that through the added effect of matching habitat choice on characteristic length and speed. The enhancement of population densities and invasion wave speeds in advective environments (Fig. 1G,H,J) suggests a complementary mechanism for the resolution of the so-called drift paradox, on top of sufficiently high random dispersal (Speirs & Gurney 2001, Nisbet et al. 2010).

Characteristic lengths, speeds and times additionally allow for recasting problems in terms of dimensionless ratios. This has value beyond simplifying the analytical or numerical solution of particular models, as it facilitates the use of *dimensional analysis* in interpretation of data and design of micro- and mesocosm experiments. Dimensional analysis has been applied successfully in engineering, geosciences, and many other scientific fields for decades.

Although there has been some discussion and application (Okubo & Levin 1989, Charnov 1993, Gaylord & Gaines 2000, Petersen & Hastings 2001, Petersen & Englund 2005, Simpson et al. 2008, Dawson & Hamner 2008, Nisbet et al. 2010, Legendre & Legendre 2012), it remains largely underutilized in ecology. By identifying a set of dimensionless ratios (such as *φ* and *Q* in this study; Eq.[5] and Eq.[7]) and a set of relationships among them, obtained from theory or empirically, one can quickly make educated guesses, without solving any sophisticated and complex models, about how a system should respond under different scenarios, beyond those directly examined. For example, the adaptation lag, Eq.[4], can be recast as 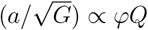, a relationship that combines three dimensionless quantities, where local maladaptaion is now measured in additive genetic standard deviations, 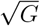. This allows for quick guesses of how local maladaptation (or biotic lag) may change under different scenarios, such as increased velocity of climate change, advective flow poleward or equatorward, or different demographic dynamics that may influence the strength of gene swamping, *φ*.

Recent microcosm experiments with swarming microbes have highlighted the distinction between pulled and pushed invasion wave fronts (Gandhi et al. 2016, Deforet et al. 2017, Birzu et al. 2019, Erm & Phillips 2020). Pulled waves are the classic invasion fronts obtained by random dispersal and negatively density-dependent growth (e.g., logistic; Shigesada & Kawasaki 1997, Murray 2002). Here, I additionally identified an opposite pulling, due to the eco-evo coupling of demography and evolution, mediated by asymmetric gene flow (May et al. 1975, Nagylaki 1978, Filin 2007), i.e., gene swamping (Lenormand 2002). In homogeneous environments, advective dispersal simply adds to the invasion speed. But when a species needs to continuously adapt to an environmental gradient in order to invade, advective dispersal intensifies the upstream pulling of gene swamping, countering the downstream pulling by dispersal and density-independent population growth at the advancing front. This slows down the evolutionary speed of species invasion (García-Ramos & Rodriguez 2002, Filin et al. 2008). In the extreme case of steep enough environmental gradients, the invasion front changes direction, and the entire range gradually, and counter-intuitively, shifts upstream.

With climate change, the effect is reversed. Pinned ranges shift downstream in the same direction as the velocity of climate change (Pease et al. 1989, Polechová et al. 2009). With both advective dispersal, *u*, and climate change, *u*_*θ*_, dynamics depends on the relative velocity, *u* − *u*_*θ*_. (Eq.[4], Eq.[5], Fig. 1E,F). Front reversals have been identified also by Potapov & Lewis (2004), in a demographic model of two competing species, under climate change. This symmetry between advective dispersal and climate change suggests some interesting directions for space-for-time substitution studies (Lovell et al. 2023), for example, by studying advective environments and the organisms that historically adapted to them as models for current and future responses to climate change elsewhere.

Gene swamping has fallen out of favor as an explanation for range limits, on both theoretical grounds (Barton 2001, Filin 2007, Polechová et al. 2009, Matz et al. 2018), and due to lack of empirical support (Angert et al. 2020, Kottler et al. 2021). (But see, recently, Hancock et al. 2024.) With the identification of a multitude of other eco-evo processes (gene surfing, expansion load, spatial sorting, etc.), gene swamping has been almost forgotten, to the point that a recent review does not even mention the term (Miller et al. 2020; cf. Lenormand 2002). In addition to causing swamping, gene flow also inflates the additive genetic variance of local populations (Slatkin 1978, Barton 2001), thus, enhancing local adaptation. In such cases, (deterministic) eco-evo models predict unlimited ranges at equilibrium (Barton 2001, Filin 2007). However, equilibrium may take a long time to achieve and, in an increasingly disturbed natural world, may be irrelevant altogether. Gene swamping is an inherent component of dispersal and gene flow in heterogeneous environments (Eqs.[1] and [13]; also Bertola et al. 2020, Hancock et al. 2024). As I have shown in this study, it has important influences on evolutionary invasion dynamics (García-Ramos & Rodriguez 2002, Filin 2007, Filin et al. 2008), particularly in cases of directional flow, advective dispersal and environmental change.

In summary, through the study of advective dispersal, I established here links among disparate modeling approaches: averaging and tracking of environmental heterogeneity and spatially varying selection (Slatkin 1973, Roughgarden 1974, May et al. 1975, Slatkin 1978, Anderson et al. 2005, Diehl et al. 2008), eco-evolutionary range and invasion dynamics along environmental gradients (Kirkpatrick & Barton 1997, García-Ramos & Rodriguez 2002, Hare et al. 2005, Filin 2007, Filin et al. 2008, Polechová et al. 2009), the drift paradox and population persistence in advective environments (Possingham & Roughgarden 1990, Gaylord & Gaines 2000, Speirs & Gurney 2001, Lutscher et al. 2006, Nisbet et al. 2010, Vasilyeva & Lutscher 2012, Meyer et al. 2024), and adaptation, persistence and critical domain size under climate change (Lande & Shannon 1996, Potapov & Lewis 2004, Berestycki et al. 2009). The symmetry between velocity of advective dispersal and that of climate change leads to unique challenges to species persistence in flow environments under global environmental change (Sorte 2013). However, due to this symmetry, populations that have historically adapted under conditions of directional flow can, additionally, function as models for persistence and adaptation under climate change in general. Such populations and species may also be preadapted to better cope with the challenges of climate change, as the eco-evo effects of advective flow are already integrated into their life-history (Müller 1982, Possingham & Roughgarden 1990, Speirs & Gurney 2001, Poulin et al. 2002, Shanks & Eckert 2005, Meyer et al. 2024). The similarity in tracking and averaging of environmental heterogeneity, in both demographic and evolutionary dynamics, points at possible application of dimensional analysis to eco-evo and climate change problems. Through demographic and evolutionary length and velocity scales, one can identify dimensionless metrics, to better understand and classify eco-evo scenarios, to inform micro- and mesocosm experiments (Petersen & Hastings 2001, Petersen & Englund 2005, Simpson et al. 2008, Gandhi et al. 2016) and space-for-time substitutions (Lovell et al. 2023), and to facilitate comparisons of different populations, species, traits and locales. A systematic application of dimensional analysis in theoretical and empirical eco-evo and climate change research has potential to provide complementary tools and insights to those of multivariate statistics.

## Appendix

As typical in modeling of transport phenomena (Beek & Muttzall 1975, Stanislav 1982, Kumaran 2023), local rate of change of a balanced quantity (e.g., population density) is the sum of of transport terms, expressing the local balance of fluxes across space (obeying some conservation law, such as of mass), and source/sink terms (hereafter *source* terms; also reaction or forcing terms) that operate locally and describe local production (or consumption; e.g., local population growh, birth rate and mortality). The following is a modified version of the derivations in Filin (2007), applying this transport vs. source distinction. I first deal with the transport terms of the dynamics, as those are the main focus of this article. The source terms are, therefore, initially abstracted away through general *S*^(*i*)^ notation, to be defined later.

A trait density function, *ρ*(*z*, ***x***, *t*) describes numbers of individuals having phenotypic value between *z* and *z* + *dz* per unit distance or area, *ρ*(*z*, ***x***, *t*)*dz*. Consequently, population density is

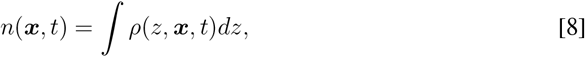

where integration is over all trait values. Local trait frequency is then given by *f* (*z*, ***x***, *t*) = *ρ/n*, or *ρ* = *nf*. The dynamics of *ρ* is described by a diffusion-advection-reaction equation,

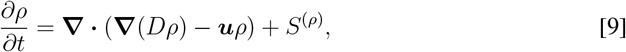

where *S*^(*ρ*)^ encapsulates the local nonconservative source processes — birth, mortality, seleciton, mutation, recombination, etc., depdending on context. The operator **∇** represents spatial derivatives, i.e., with respect to spatial position, ***x***. In Eq.[9], **∇** describes how spatial variation in net fluxes of random and advective dispersal add to local population growth, on top of source terms. Diffusive transport (random dispersal) is denoted by *D*, while advective (or convective) transport (directional dispersal) by a velocity vector ***u***. The expression **∇**(*D*(***x***, *z, t*)*ρ*(*z*, ***x***, *t*)) − ***u***(***x***, *z, t*)*ρ*(*z*, ***x***, *t*), then, represents the net flux of individuals of trait value *z* at location ***x*** at time *t*.

Rewriting Eq.[9] in terms of *n* and *f*, I obtain

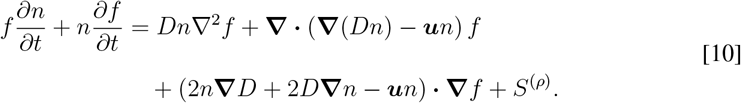

By isolating *∂f* /*∂t*, I can further derive the following equation for the dynamics of trait frequency distribution, *f*,

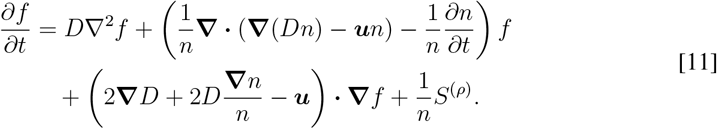

Furthermore, by integrating Eq.[9] over all trait values (over *z*) to get the equation for *∂n*/*∂t*, I finally obtain the coupled dynamics equations for the spatially-varying population density and trait frequency distribution (probability density function) are given by

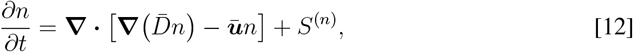

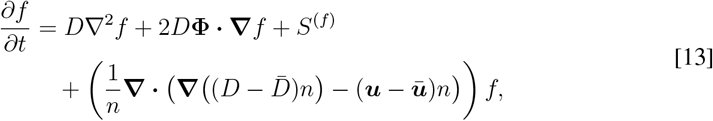

where *S*^(*n*)^ and *S*^(*f*)^ are general source terms for population density and trait frequency, respectively, 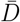 and 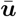 are mean diffusive and advective transport in the local population (population means of *D* and *u*), and the gene flow asymmetry vector, **Φ**, is as defined in Methods (Eq.[1]).

Next, I simplify the trait dynamics by following only that of the mean trait value, given by 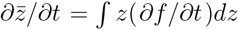, where *∂f* /*∂t* is given by Eq.[13]. The eco-evolutionary dynamics equation 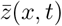 is then, given constant diffusive and advective dispersal (*D* = *const, u* = *const*),

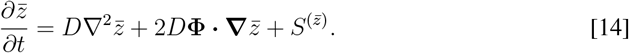

Given the linearly varying environmental gradient, as described in Methods, Eq.[14] can be rewritten as an equation for local (mal)adaptation, 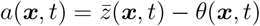 (i.e., adaptation lag, or biotic lag),

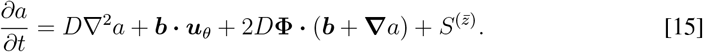

With trait-dependent habitat choice, the sorting term, 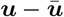, in Eq.[13] does not vanish and, therefore, modifies the dynamics of 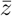 (or *a*), when evaluating the integral 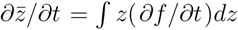. As also derived in Filin (2007), given a stabilizing selection of intensity *s*, such that fitness contains a quadratically trait-dependent term, 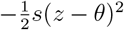, these extra habitat choice and sorting terms add up to 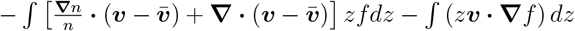, where 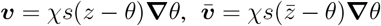, and 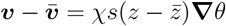. The resulting dynamics of mean trait is then given by

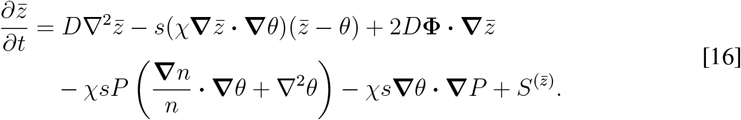

Here, **Φ** = **∇** ln(*n*) − ***u****/*2*D*, ***u*** being the fixed (trait-independent) advective dispersal component. *P* denotes the trait (phenotypic) variance. Again, given the directionally changing linear environmental gradient, the equation for adaptation lag, *a*(***x***, *t*), is

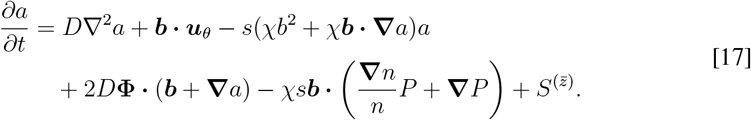

At this point, it is important to note that the above expressions were obtained by just analyzing the transport terms for dynamics of population density, trait distribution, and mean trait or adaptation lag. No assumption was made regarding the source terms, *S*^(*ρ*)^, *S*^(*n*)^, *S*^(*f*)^ and 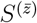, or the shape of the trait distribution, *f*. In particular, the sorting term −*s*(*χb*^2^)*a* (a “restoring force”), is equivalent to stabilizing selection, where 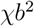 plays a role akin to additive genetic variance (see below). This equivalence, again, does not depend on the shape of the trait distribution.

Finally, the specific model, explored in the numerical examples, makes the simplifying assumptions of one-dimensional space, ***x*** = *x* (dropping the vector notation). Additionally, the source terms for population density, *S*^(*n*)^, and mean trait, 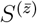, describe logistic population growth and stabilizing selection, such that Eqs.[12] and [15] become

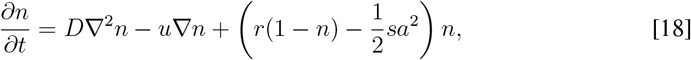

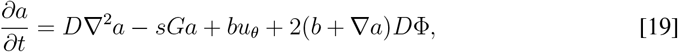

where now 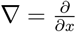. Without loss of generality, the genetic load term, 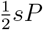, has been absorbed into the definition of *r* and the carrying capacity (García-Ramos & Rodriguez 2002), and population density is scaled such that carrying capacity is 1. The phenotypic and additive genetic variances, *P* and *G*, are taken to be fixed parameters, or, at least, varying much more slowly than *n* and 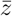. (Eqs.[18] and [19] are, up to differences in notation [see SI] and the extra advection terms, the same as previously presented for eco-evo dynamics along environmental gradients; Pease et al. 1989, Kirkpatrick & Barton 1997, Case & Taper 2000, García-Ramos & Rodriguez 2002, Hare et al. 2005, Polechová et al. 2009.)

Similarly, the equations for eco-evolutionary dynamics, with trait-dependent habitat choice, become

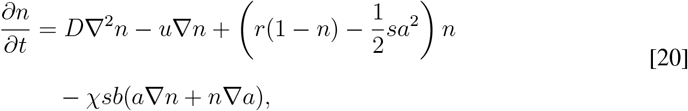

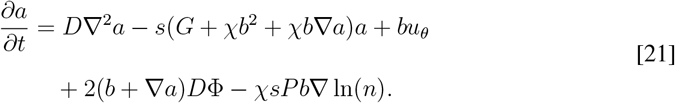

Compare the latter with Eq.[17], now fully demonstrating the equivalence of *χb*^2^ to additive genetic variance, *G*, and the “restoring force” role of both sorting by habitat choice and stabilizing selection.

### THE EVOLUTIONARY AVERAGING KERNEL

In constant environments, *θ* = *θ*(***x***) or *u*_*θ*_ = 0, and recalling that by definition 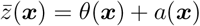, one can obtain from Eq.[19] an expression for the mean phenotype at equilibrium,

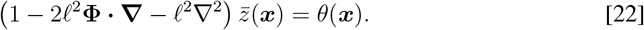

Dropping the vector notation for the one-dimensional case (***x*** = *x*), it is straightforward to show, by Fourier transform or Green’s function, that this expression defines an averaging scheme over *θ*(*x*), or low-pass filter (Slatkin 1978, Filin 2007). The averaging kernel is given by two separate decaying exponentials for positive and negative displacements from the focal locality, *x* of 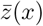, corresponding to downstream and upstream directions,

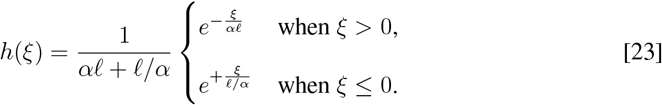

The parameter *α* represents the contraction downstream and expansion upstream of the averaging kernel, due to asymmetric gene flow, 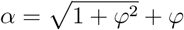, where *φ* = *ℓ*Φ. The mean phenotype at equilibrium is then given by 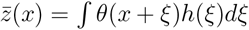, a low-pass-filtered optimum.

With climate change, *u*_*θ*_ *>* 0, a transformation of the space variable *x* to *x* − *u*_*θ*_*t* restores a constant (time-invariant) environmental gradient. In this case, Φ (Eq.[6]) includes advection velocity of *u* − *u*_*θ*_, which leads to *φ* according to Eq.[7].

### APPROXIMATE EXPRESSION FOR INVASION SPEED

Under a change of space variable, *y* = *x* − *ct*, I look at a traveling wave solution of the form *n* = *n*_0_*e*^−*κy*^ and *a* = *a*_0_ − *qy, q, κ >* 0, for large *y*, i.e., at the advancing front, where population density is low, and growth is thus approximately density-independent. Substitution into Eq.[18] and Eq.[19] yields 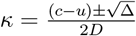, where the discriminant, Δ, is given by 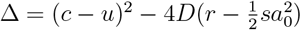. The expression for baseline maladaptation, *a*_0_, is *a*_0_ = *b*(*u*_*θ*_ − *c*)*/sG*. Substituting into the equation for the discriminant and setting Δ = 0 (Shigesada & Kawasaki 1997, Murray 2002) finally yields Eq.[5] for invasion speed, *c*.

## Acknowledgements

This work began over 20 years ago. The general phenotypic model and some other elements were originally written in a declined manuscript in 2004. I presented them at the 2004 Society for Mathematical Biology (SMB) annual meeting in Ann Arbor. I thank the SMB for granting a 2004 Landhal Travel Award. Eventually that early work ended up as two chapters in my PhD dissertation. That phase of the research was conducted in Yaron Ziv’s Spatial Ecology lab at Ben-Gurion University, Beer-Sheva, Israel. I thank Prof. Ziv for support and advice during those years. Similarly, I acknowledge a Kreitman Foundation Doctoral Fellowship, granted 2003-2007. I also acknowledge helpful comments by Nick Barton, Mark Blows, Jonathan Losos and an anonymous reviewer, in 2004. I thank Bob Holt for encouraging me to get this work published already back in 2009. It took a while. Finally, I thank Jitka Polechováfor discussion and advice during a recent visit to Vienna.

