## Supplementray material for "Symmetry and similarity in eco-evo: advective environments, dispersal, and climate change"

### Supplementary information for “Symmetry and similarity in eco-evo: advective environments, dispersal, and climate change”

Ido Filin [1] *

[1] Kangasala, Finland

#### Contents

1 A guide to notation 2

2 Numerical procedure 2

3 Supplementary references 4

DRAFT

#### A guide to notation

Population genetic studies (Lande 1982, Bürger & Lynch 1995, Kirkpatrick & Barton 1997, Polechová et al. 2009) usually describe random dispersal using the second infinitesimal moment (Karlin & Taylor 1981) of the dispersal diffusion process, $\sigma^2$, instead of the diffusion coefficient, $D$. But the two are related by $\sigma^2 = 2D$.

The fitness function in population-genetic studies is usually given by $r(1-n) - \frac{(z-\theta)^2}{2V_s}$, where $V_s$ is inversely related to strength of stabilizing selection. It is related to $s$ of this study by $V_s = 1/s$. Filin (2007) used $r(1-n) - s(z-\theta)^2$, which produces an extra factor of 2 in the dynamics of mean trait or maladpataion. In this study I opted for the expression $r(1-n) - \frac{1}{2}s(z-\theta)^2$, which removes this extra factor. All three forms are equivalent.

The dimensionless ratio, $Q$, given by $Q^2 = 2Db^2/sG^2$, is related to the dimensionless variables $A$ and $B$ of Kirkpatrick & Barton (1997) through $Q^2 = 2B^2/A^2$. This ratio also appears in Miller (2019), in relation to critical velocity. The dimensionless maladaptation expression, $a^*$, of Polechová et al. (2009) is related to the dimensionless ratios in this study by $a^* = \varphi Q\sqrt{2A}$.

#### Numerical procedure

Selecting space, time and trait scales, such that $D = r = s = 1$, the dynamics equations (Eqs.[18] and [19] in the Appendix) are transformed to

$$\frac{\partial n}{\partial t} = \nabla^2 n - u\nabla n + \left(1 - n - \frac{1}{2}a^2\right) n, \tag{S1}$$

$$\frac{\partial a}{\partial t} = \nabla^2 a - Ga + bu_\theta + 2(b + \nabla a)\Phi, \tag{S2}$$

Under such rescaling, eqn S1 and eqn S2 are the same as the dimensionless equations of García-Ramos & Rodriguez (2002), with extra terms for advective dispersal and climate change. The remaining two parameters, $G$ and $b$, are respectively equivalent to $A$ and $B$ of Kirkpatrick & Barton (1997), García-Ramos & Rodriguez (2002), Polechová et al. (2009).

Using rendering buffers of size 32x4096 single-precision (32-bit) floating-point pixels in video memory, I am able, at each simulation run, to study the effect of 32 different values of $b$ on eco-evo dynamics. Space is represented by a one-dimensional spatial mesh of 4096 discrete locations, separated by $\delta x$, ranging between $0.0125$ and $0.025$. The time step was chosen accordingly as $\delta t = 0.24(\delta x)^2$ (Smith 1985). I used convective boundary conditions, such that $\ln(n)$ and $a$, and their spatial derivatives, were extrapolated to positions $x_{-1}$ and $x_N$ ($N = 4096$) at each time iteration.

DRAFT

DRAFT
